# URMC-099 Prophylaxis Prevents Hippocampal Vascular Vulnerability and Synaptic Damage in an Orthopedic Model of Delirium Superimposed on Dementia

**DOI:** 10.1101/2021.10.04.463042

**Authors:** Patrick Miller-Rhodes, Herman Li, Ravikanth Velagapudi, Niccolò Terrando, Harris A Gelbard

## Abstract

Systemic perturbations can drive a neuroimmune cascade after surgical trauma, including affecting the blood-brain barrier (BBB), activating microglia, and contributing to cognitive deficits such as delirium. Delirium superimposed on dementia (DSD) is a particularly debilitating complication that renders the brain further vulnerable to neuroinflammation and neurodegeneration, albeit these molecular mechanisms remain poorly understood. Here we have used an orthopedic model of tibial fracture/fixation in APPSwDI/mNos2^-/-^ AD (CVN-AD) mice to investigate relevant pathogenetic mechanisms underlying DSD. We conducted the present study in 6 months-old CVN-AD mice, an age at which we speculated amyloid-β pathology had not saturated BBB and neuroimmune functioning. We found that URMC-099, our brain-penetrant anti-inflammatory neuroprotective drug, prevented inflammatory endothelial activation, breakdown of the BBB, synapse loss, and microglial activation in our DSD model. Taken together, our data link post-surgical endothelial activation, microglial MafB immunoreactivity, and synapse loss as key substrates for DSD, all of which can be prevented by URMC-099.

## Introduction

Delirium is a common neurologic complication frequently encountered after major surgery, such as cardiac and orthopedic, which can impact neurodegenerative processes and contribute to significant mortality and morbidity in older and frail patients [1]. Indeed, patients with Alzheimer’s disease (AD) often require orthopedic surgery to repair a broken limb and become particularly vulnerable to complications, including postoperative delirium (POD), an acute state of cognitive impairment that develops in the hospital setting [2–4]. POD is associated with an increased one-year mortality and a worsening of cognitive trajectories in patients with pre-existing neurodegeneration [5, 6]. Despite the prevalence of POD in this population, the pathophysiology of delirium superimposed on dementia (DSD) remains poorly understood and without disease-modifying or prophylactic therapeutic interventions.

Vascular etiologies for cognitive impairment and dementia are a growing research interest, yet the impact of systemic stressors on the neurovascular unit (NVU) is far less established, especially in the context of perioperative complications. To better understand the putative impact of surgery on neurovascular and immunological sequelae we have developed a model of DSD following tibial fracture and fixation in CVN-AD mice [7]. In the absence of surgical manipulation, the CVN-AD model displays age-dependent effects including amyloid-β accumulation [8], cerebral amyloid angiopathy (CAA), tau phosphorylation [8–10], microgliosis [8], astrogliosis [9], NVU dysfunction [11], increased expression of complement components [9], and neuronal degeneration [10]. In our prior work we have established that tibial fracture/fixation in this mouse accelerates neurodegenerative pathology, especially amyloid-β accumulation in the dentate gyrus (DG) region of hippocampus, which is accompanied by delirium-like behavior as defined by inattention deficits in the 5-choice serial reaction time task. These pathologic and behavioral changes were particularly evident in 12 months-old CVNs and accompanied by greater NVU pathology, dramatic loss of AQP4, and concomitant fibrinogen deposition [7]. Since vascular factors, including vascular cell adhesion molecule type 1 (VCAM-1), have been identified in a proteomics screen as key gateway proteins associated with pathologic aging [12], we sought to investigate whether surgery impacted VCAM-1 expression and whether treatment with URMC-099 in CVN-AD could prevent neurovascular pathology in DSD.

Here, we test the hypothesis that URMC-099 prevents damage to the blood-brain barrier (BBB) and hippocampal synapses resulting from orthopedic surgery in 6 and 12 month-old mice, the former representing an early stage in AD progression and the latter late-stage disease.

## Materials and methods

### URMC-099

URMC-099 (M.W. 421) was synthesized as originally described in Goodfellow et al. [13]. URMC-099 drug solutions were prepared by dissolving 20 mg of URMC-099 in 0.5mL sterile dimethyl sulfoxide (DMSO; D8779, Sigma-Aldrich, St. Louis, MO). We then added 4 mL polyethylene glycol 400 (PEG400; 91893-250-F, Sigma-Aldrich) and 5.5mL sterile saline (National Drug Code NDC0409-4888-10) for a final concentration was 2mg/mL URMC-099. The vehicle was the same solution minus URMC-099. Mice were administered (i.p.) three injections at 12 h intervals prior to orthopedic surgery. Drug solutions were coded such that experimenters were blinded to the experimental conditions for the duration of the experiments.

### Cell culture

Bend.3 cells [14] (ATCC CRL-2299; BALB/c origin; generously provided by Dr. Ning Quan) were cultured in glutamate-free Dulbecco’s Modified Eagle’s Medium (DMEM) supplemented with GlutaMAX (1% v/v) and fetal bovine serum (FBS; 10% v/v) and maintained in a humidified 37° C incubator (5% CO2). For immunocytochemistry (ICC), bend.3 cells were seeded at a density of 100,000 cells per well on plasma-cleaned, poly-D-lysine-coated coverslips in a 12-well plate and allowed to adhere overnight. Subsequently, bend.3 cells were serum starved for 30 minutes, followed by addition of URMC-099 (300 nM in DMSO). Thirty minutes after URMC-099 pre-treatment, PBS or IL-1β (10 ng/mL, a dose informed by [12]) was added as indicated. Bend.3 cells were fixed 16 hr post-IL-1β stimulation for ICC. For quantitative reverse-transcriptase polymerase chain reaction (qRT-PCR), bend.3 cells were treated exactly as above, except cells were seeded at 200,000 cells per 12-well and lysates were collected at 6 hr post-IL-1β stimulation. Because biological reproducibility cannot be measured in cell lines, all cell line experiments were performed once, with all values and statistics computed across replicate wells. Finally, viability assays were not formally assessed during these experiments because of the lack of morphologic criteria for cell death in all culture wells.

### Animals and orthopedic surgery

APPSwDI/mNos2^-/-^ AD mice (CVN-AD; kindly provided by Dr. Carol Colton, Duke University) were bred and maintained by the Terrando laboratory at Duke University. The procedures followed to provide tissues from APPSwDI/mNos2^-/-^ AD mice were performed in strict compliance with animal protocols approved by the Institutional Animal Care and Use Committees (IACUC) of Duke University (A249-17-11). Mice were housed under a 12-hr light/dark cycle with free access to water and regular chow. URMC-099 prophylaxis and orthopedic surgery were performed as described in Miller-Rhodes et al., 2019 [15] on 6-month and 12-month of APPSwDI/mNos2^-/-^ mice under isoflurane (Patterson Veterinary, Greeley, CO) anesthesia and analgesia (buprenorphine, 0.1 mg/kg subcutaneously; ZooPharm, Laramie, WY). Briefly, a small incision was performed on the shaft of the tibia followed by muscle disassociation and stripping of the periosteum. Pinning of the tibia was performed through the insertion of a 0.38mm stainless steel rod and osteotomy performed on the upper crest of the bone. All mice recovered from surgery and were included in the study.

### Immunocytochemistry

Immunocytochemistry was performed per Glynn et al., 2006’s protocol [16] with minor modifications. Briefly, following fixation in 4% PFA/4% sucrose solution for 15 min at 4°C, cells were washed with 100 nM glycine (in 1X PBS), followed by two more washes in 1X PBS. Cells were permeabilized in 0.25% Triton-X for 5 min (followed by 2 PBS washes), blocked in 10% bovine serum albumin (BSA) in 1X PBS, and incubated with primary antibodies in 3% BSA for 1 hr at room temperature. Primary antibodies included rat monoclonal anti-VCAM1 (Abcam, ab19569, clone M/K-2; 1:250) and Armenian hamster anti-PECAM-1 (CD31; Millipore Sigma, cat no. MAB1398Z, clone 2H8; 1:250). Cells were washed again three times with 1X PBS and incubated with secondary antibodies in 3% BSA for 45 min at room temperature. Cells were washed a final three times with 1X PBS, dipped in ddH20 to remove excess salt, and mounted on microscope slides with Prolong Diamond anti-fade reagent (ThermoFisher, cat no. P36961).

### Quantitative reverse transcriptase polymerase chain reaction (qRT-PCR)

RNA was extracted using a spin column RNA isolation kit (Machery-Nagel). On-column DNase treatment was used to remove contaminating genomic DNA. Total RNA preparations were then used directly for qPCR (200 ng total RNA per reaction) using the Luna Universal One-Step RT-qPCR Kit (NEB, E3006) and target-specific primer/probe sets (ThermoFisher) or frozen at −80° C until use. The qPCR primer/probe sets we used include the endogenous control *Ipo8* (Mm01255158_m1), *Vcam1* (Mm01320970_m1), *Spp1* (Mm00436767_m1), *Timp1* (Mm01341361_m1), and *Col1a1* (Mm00801666_g1). The ΔΔCt method was used to analyze the results.

### Immunohistochemistry

Twenty-four hours after orthopedic surgery, mice were deeply anesthetized with isoflurane and transcardially perfused with PBS and subsequently 4% PFA. Brains were carefully extracted and post-fixed for 24 hr in 4% PFA. Brains were transferred to conical tubes containing PBS, packed with ice-packs, and shipped to the Gelbard laboratory at University of Rochester Medical Center. Free-floating coronal brain sections (40μm thickness) were cut using a vibratome and stored in cryoprotectant (30% PEG300, 30% glycerol, 20% 0.1 M phosphate buffer, and 20% ddH2O) at −20° C. For immunohistochemistry, cryopreserved sections were washed three times in 1X PBS followed by another wash in 0.1 M glycine in 1X PBS (to reduce autofluorescence). Sections were subsequently incubated in blocking buffer (1.5% BSA, 3% normal goat serum, 0.5% Triton-X, and 1.8% NaCl in 1X PBS) containing primary antibodies (VCAM1 Millipore Sigma MAB1398Z 1:200; Tmem119 SySy 1:500; Iba1 Wako PTR2404 1:1000; MafB Atlas HPA005653 1:500; CD68 Serotec MCA1957GA 1:1000; Homer1 SySy 160006 1:500; PSD95 NeuroMab 75-028 1:500; Piccolo SySy 142104 1:500; C1q Abcam AB71940 1:500; CD31 Millipore Sigma MAB1398Z 1:250; Fibrinogen Dako A0080 1:200; H3Cit Abcam ab5103 1:500) in blocking buffer (1.5% BSA, 3% normal goat serum, 0.5% Triton-X, and 1.8% NaCl in 1X PBS) overnight at room temperature. Then sections were washed thrice in 1X PBS containing 1.8% NaCl before incubating in Alexa Fluor conjugated secondary antibodies (ThermoFisher), again overnight. Finally, sections were washed three times with 1X PBS+1.8% NaCl, mounted on glass slides, and cover-slipped with Prolong Diamond Antifade Reagent (Invitrogen P36961). When used, the UV-excitable fibrillar amyloid-β dye Amylo-Glo (Biosensis TR-300-AG; 1:100) or Alexa-633 Hydrazide (Thermo Fisher A30634; 1:1000) was diluted in 1X PBS+1.8% NaCl and applied to sections for 10 min following the first two post-secondary washes; an additional two washes (1X PBS+1.8% NaCl) were performed to rinse the sections of excess Amylo-Glo dye or Alexa-633 Hydrazide.

### Grid pseudo-confocal microscopy and image analysis

Slides were coded throughout imaging and analysis to reduce experimenter bias. We used an Olympus BX-51 microscope equipped with Quioptic Optigrid hardware (for optical sectioning) and a Hamamatsu ORCA-ER camera. Images were acquired using Volocity 3DM software (Quorum Technologies). Within each set, light intensity and exposure settings were kept constant. For ICC, three images were acquired for each experimental replicate. For IHC, six images were acquired for each biological replicate.

Synaptic punctae were imaged at 60X magnification (10μm depth, 0.3μm step size). To reduce variability due to differences in vascular coverage (i.e., areas without synaptic punctae), we selected 300 x 300 pixel ROIs from each XYZ-stack for analysis. Other markers were imaged at 20X and 40X magnification. Immunostained objects were identified and quantified using custom Volocity workflows. More specifically, for synaptic punctae, the colocalization of spots and objects were used to identify punctae. The “Find Spots” function was used to identify individual puncta based on local intensity maxima and “Find Objects” function was used to identify punctae with a minimum intensity threshold; all other quantified objects were identified using the “Find Objects” function. Arterial VCAM-1 vessels were defined as VCAM-1 objects touching Alexa-633 objects. Tmem119^lo^ and Tmem119^hi^ Iba1-positive cells were determined by intensity thresholding (by standard deviation; SD = 1.2). Microglial MafB intensity was quantified as the sum intensity of MafB objects contained within Iba1-positive objects and normalized to the number of microglia in each field of view. Non-microglial MafB was quantified as the sum intensity of MafB objects *not* contained within Iba1-positive objects and normalized to the number of MafB+ nuclei per field. Microglial CD68 was quantified as the sum intensity of CD68 objects contained with Iba1-positive objects.

### Statistics

All qPCR statistics were performed on ΔCt values and then represented graphically as fold change (2^-ΔΔct^). One- and two-way analysis of variance (ANOVA) with Holm-Sidak’s or Tukey’s multiple comparison tests were used to analyze the ICC and IHC data as indicated in each figure legend. Pearson’s correlation was used to compute correlations between VCAM-1 immunoreactivity and fibrillar amyloid-β volume. All data are presented as mean ± standard error (SEM) with significance at P < 0.05.

## Results

### Effect of URMC-099 on brain endothelial cell activation in vitro

Although URMC-099’s pharmacokinetic properties allow it to readily cross the BBB and achieve therapeutic concentrations in the CNS to act directly on microglia, its therapeutic efficacy in *in vivo* models of systemic inflammation may be attributable, at least in part, to actions on the cerebrovascular endothelium. To test whether URMC-099 can inhibit the activation of brain endothelial cells (BECs) under inflammatory conditions, we analyzed VCAM-1 and PECAM-1 (CD31) immunoreactivity (IR) on bend.3 cells, a murine cerebral microvascular endothelial cell line [14], following stimulation with the pro-inflammatory cytokine IL-1β (10 ng/mL, as previously used in [12]) alone or in combination with URMC-099 (300 nM). Twenty-four hours following IL-1β stimulation, bend.3 cells upregulated VCAM-1 (p = 0.0003), but not PECAM-1 immunoreactivity (IR) (**Figure 1A-C**). URMC-099 pretreatment virtually abolished the IL-1β-induced increase in VCAM-1 IR (p < 0.001; **Figure 1A, C**). Additionally, it also reduced the basal IR of PECAM-1 by nearly half (p = 0.001; **Figure 1A, B**). We next characterized the effect of URMC-099 on the transcriptional response of bend.3 cells to IL-1β. In addition to *Vcam1*, we assayed the expression of *Spp1, Timp1*, and *Col1a1* — all of which are upregulated in activated BECs *in vivo* in murine disease models of core BBB dysfunction [17]. We focused on *Spp1, Timp1* and *Col1a1* because of their respective roles in BBB repair following injury [18], preserving BBB integrity [19] and functional integrity of extracellular matrix proteins [20], respectively. IL-1β upregulated *Vcam1* expression 59.16-fold (p < 0.0001) and elicited smaller (<2-fold) increases in *Spp1* (p = 0.0002) and *Timp1* (p = 0.0004) mRNA (**Figure 1D**). Of these genes, URMC-099 pretreatment significantly reduced *Vcam1* (p < 0.0001) and *Spp1* (p = 0.0008) mRNA levels (**Figure 1D**). In addition, IL-1β decreased the expression of *Col1a1* by 25% versus control (p = 0.0077), an effect that was not modulated by URMC-099 pretreatment (**Figure 1D**). Together, these experiments demonstrate that URMC-099 exerts anti-inflammatory effects on BEC activation *in vitro*.

**Figure 1.**
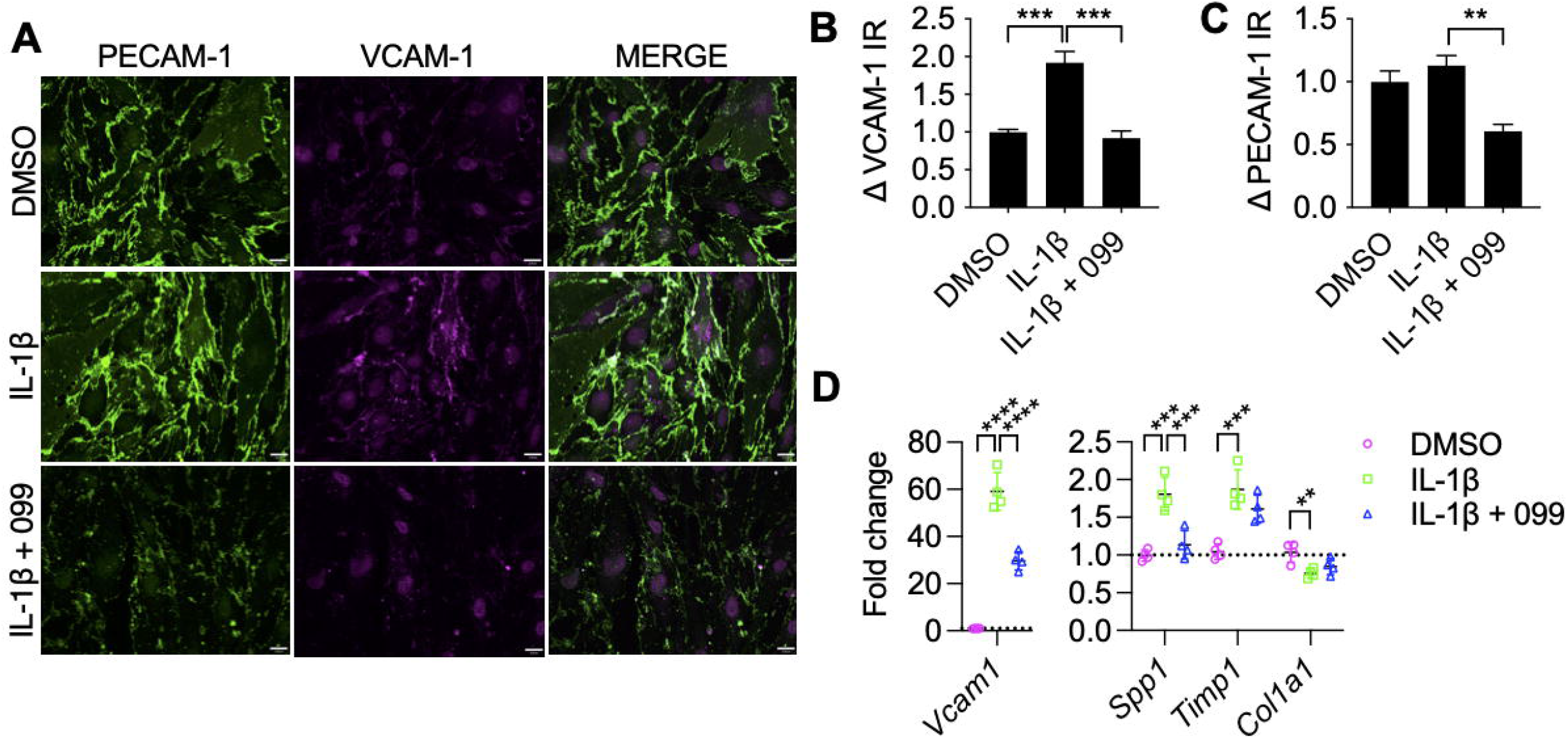
URMC-099 inhibits BEC activation *in vitro*. Serum-deprived Bend.3 cells were pretreated with URMC-099 (099) or DMSO for 30 min, after which cells were stimulated with IL-1β (10 ng/mL). (A) Representative images of VCAM-1 and PECAM-1 staining in bend.3 cells; scale bar = 15 μm. (B) Quantification of PECAM-1 IR, represented as fold change. (C) Quantification of VCAM-1 IR, represented as fold change. (D) qPCR analysis of bend.3 transcriptional response to IL-1B and URMC-099 treatment. Values presented as mean ± SEM (N = 4 cell culture wells). ** P < 0.01, *** P < 0.001, **** P < 0.0001 for indicated comparisons; one-way ANOVA with Dunnett’s multiple comparison test (B-D).

### URMC-099 prophylaxis inhibits cerebrovascular VCAM-1 IR after orthopedic surgery in CVN-AD mice

Based on our *in vitro* BEC results, we hypothesized that URMC-099 prophylaxis would inhibit cerebrovascular inflammation in the SLM of CVN-AD mice following surgery. Orthopedic surgery elicited a 1.7-fold increase in total VCAM-1 volume in the SLM relative to naïve, vehicle-treated controls (18542.5 ± 2318.7 μm^3^ *vs* vehicle-treated controls 10840.8 ± 2825.2 μm^3^; p < 0.0001; **Figure 2A-D, E**). In agreement with our *in vitro* data, URMC-099 prophylaxis decreased total VCAM-1 in the SLM (12679.7 ± 1698.5 μm^3^; p = 0.0006; **Figure 2E**). To determine whether these effects were specific to arterial or venous BECs, we used the arterial-specific dye Alexa 633 hydrazide [21] — which binds to elastin fibers present in arterioles — to distinguish between arterial and venous VCAM-1 objects in our analysis. Arterial VCAM-1 accounted for 27.3% of total VCAM-1 volume; further, there were no main effects of surgery [F (1, 20) = 3.53; p = 0.08] nor URMC-099 treatment [F (1, 20) = 0.22; p = 0.65] on arterial VCAM-1 levels (**Figure 2A’-D’, E**). In contrast, we observed significant main effects of surgery [F(1, 20) = 15.27; p = 0.0009] and URMC-099 [F(1, 20) = 14.32; p = 0.0012] on venous VCAM-1 levels, mirroring the differences between groups observed in total VCAM-1 volume. In order to eliminate the possibility of more vessels as a contributor of increased VCAM-1 volume we assessed VCAM-1 fluorescent intensity normalized to its volume. We found orthopedic surgery induced a 47% increase in VCAM-1 density compared to sham mice (5873.6 ± 425.2 A.U./μm^3^ vs sham 3993.9 ± 108.4 A.U./μm^3^; p < 0.001; **Figure 2F**). Further, pretreatment of URMC-099 prevented VCAM-1 induction in mice that received orthopedic surgery compared to vehicle treated mice (4949.8 ± 275.6 A.U./μm^3^; p < 0.05; **Figure 2F**). To rule out a potential contribution of Aβ burden on cerebrovascular VCAM-1 levels, we visualized fibrillar Aβ in the SLM using the UV dye Amylo-Glo (**Figure 2A-D**). We did not detect significant differences in fibrillar Aβ between any experimental condition (**Figure 2G**), nor was there any correlation between fibrillar Aβ levels and total VCAM-1 levels in the SLM (**Figure 2H**). Overall, these results indicate that URMC’s effect is to reduce capillary and venular VCAM-1 levels in the SLM. These data further suggest that therapeutic intervention earlier in the course of neuropathogenesis may elicit better outcomes in this DSD model. In support of this interpretation, experiments in 12 mos-old CVN mice demonstrated advanced pathology in hippocampal SLM with limited efficacy by URMC-099 in the late stages of the disease (**Figure 2I-J**).

**Figure 2.**
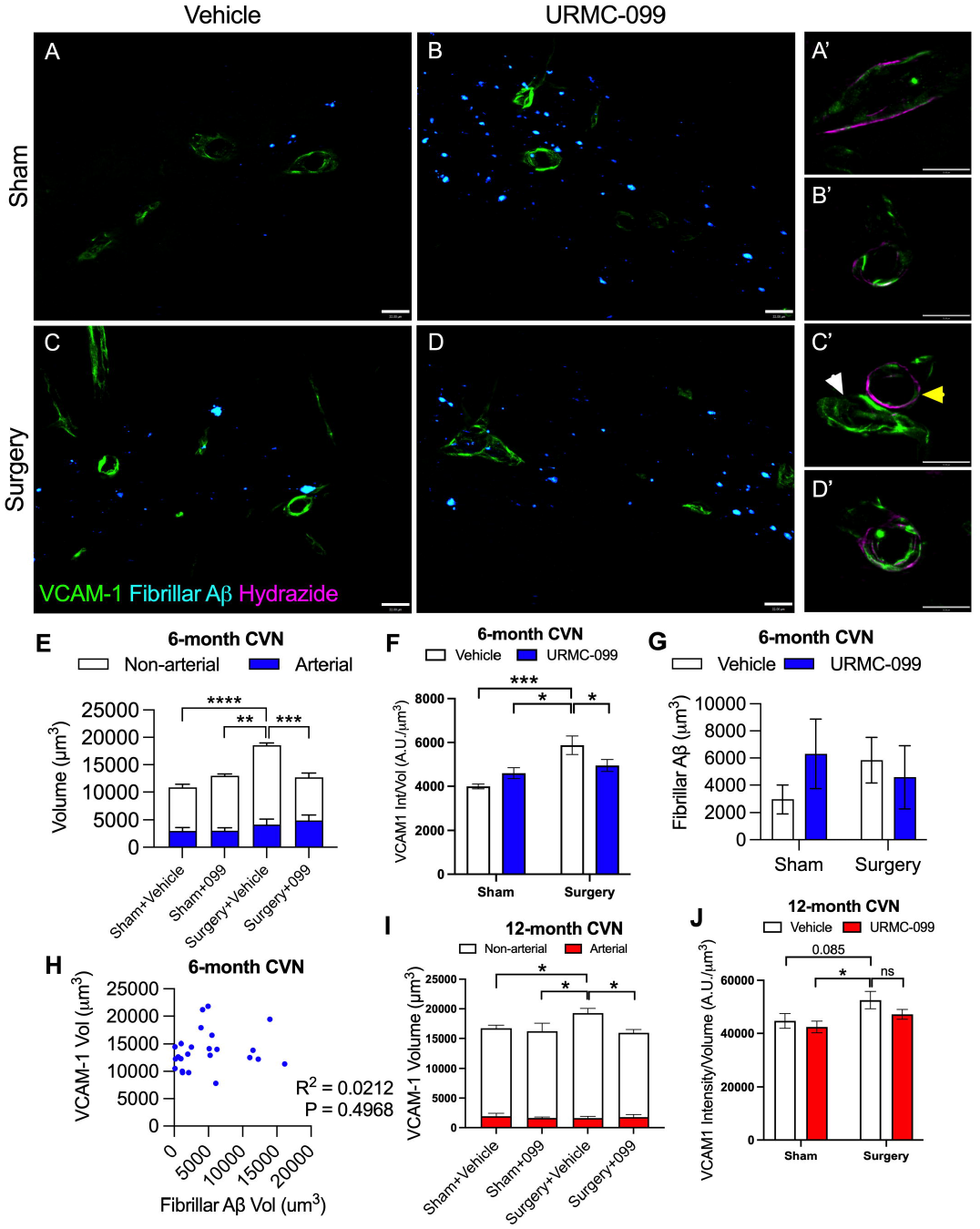
URMC-099 prophylaxis prevents the induction of VCAM-1 in the SLM of 6-month-old CVN-AD mice following surgery. Six and twelve-month-old CVN-AD mice (N=6/group) received three doses i.p. of URMC-099 (10 mg/kg) prior to undergoing sham or orthopedic surgery. Brains were harvested 24 h post-surgery for IHC. (A-D) Representative images depicting VCAM-1 (green) and fibrillar AB (cyan). (A’-D’) Representative images depicting arterial VCAM-1 vessels using the artery-specific dye Alexa 633-hydrazide (magenta); images correspond to experimental conditions represented by A-D; scale bar = 32μm. (E) Quantification of total VCAM-1 volume (full bars), venous VCAM-1 volume (white bars), and arterial VCAM-1 volume (blue bars) at 6-months age; statistical comparisons are shown for total VCAM-1. (F) Quantification of VCAM-1 intensity normalized to volume at 6-months of age. (G) Quantification of fibrillar AB. (H) Correlation between total VCAM-1 volume and fibrillar AB volume at 6-months of age. (I) Quantification of total VCAM-1 volume (full bars), venous VCAM-1 volume (white bars), and arterial VCAM-1 volume (blue bars) at 12-months age; statistical comparisons are shown for total VCAM-1. (J) Quantification of VCAM-1 intensity normalized to volume at 12-months of age. Values presented as mean ± SEM (N = 6); * P<0.05, ** P< 0.01, *** P < 0.001, **** P < 0.0001 for indicated comparisons; two-way ANOVA with Holm-Sidak’s multiple comparison test (E, F) or Pearson’s coefficient (D)

### URMC-099 prophylaxis prevents vascular neutrophil NET formation and barrier dysfunction after orthopedic surgery in CVN-AD mice

To further evaluate barrier dysfunction in CVN-AD mice after surgery we measured fibrinogen extravasation out of CD31+ vessels in the SLM at 24-hours (**Figure 3A**). Surgery induced a robust 2.8-fold increase of fibrinogen out of the vascular lumen compared to naïve vehicle-treated controls (1931.4 ± 508.4μm^3^ vs sham 687.6 ± 105.7μm^3^ vs; p < 0.05), which was effectively prevented by URMC-099 (920.110 ± 224.1μm^3^; p < 0.05; **Figure 3B**) with arrows being sites of fibrinogen leakage outside of the lumen. Extravascular fibrinogen deposition was co-localized with an induction of CD31 expression (**Figure 3C**), further highlighting the key role of neurovascular damage in this model of DSD. To assess the contribution of neutrophils to barrier dysfunction, we measured the formation of neutrophil extracellular traps (NETs) using citrullinated histone H3 (H3Cit) a hallmark of NETs. Interestingly, we found orthopedic surgery caused significant 10.2-fold increase of NETs along the vascular lumen in the SLM (822.7 ± 351.7μm^3^ vs vehicle-treated sham 80.1 ± 22.2μm^3^; p < 0.05; **Figure 3D-E**). Formation of perivascular NETs was prevented with pretreatment of URMC-099 compared to vehicle-treated surgical mice (110.2 ± 38.8μm^3^; p < 0.05; **Figure 3D-E**). Interestingly, histological assays did not show any visible Ly6G+ neutrophils in the SLM (data not shown).

**Figure 3.**
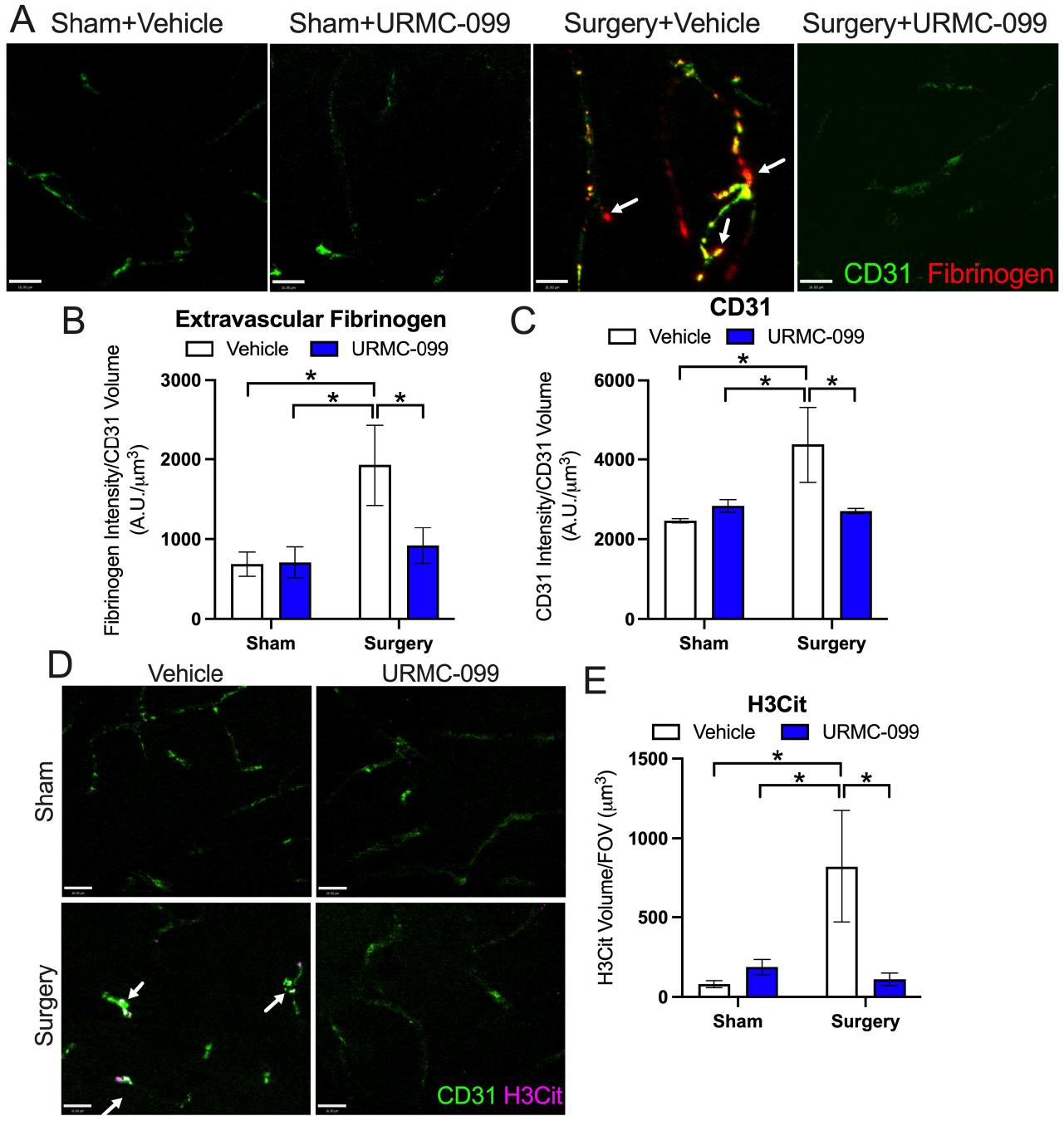
URMC-099 prevents vascular damage in the SLM of 6-month CVN-AD mice following orthopedic surgery. Six-month-old CVN-AD mice (N=6/group) received three doses i.p. of URMC-099 (10 mg/kg) prior to undergoing sham or orthopedic surgery. Brains were harvested 24 h post-surgery for IHC. (A) Representative images depicting CD31/Pecam1 (green) and fibrinogen (red) with arrows being sites of fibrinogen leak; scale bar = 16 μm. (B) Quantification of extravascular fibrinogen. (C) Quantification of CD31/Pecam1 intensity normalized to CD31/Pecam1 volume. (D) Representative images depicting CD31/Pecam-1 (green) and NET marker, H3Cit (magenta) with arrows being sites of colocalization; scale bar = 16 μm. (E) Quantification of neutrophil NET marker, H3Cit. Values presented as mean ± SEM (N = 6); * P < 0.05 for indicated comparisons; two-way ANOVA with Holm-Sidak’s multiple comparison test (C-E).

### URMC-099 prophylaxis ameliorates synapse loss following orthopedic surgery in CVN-AD mice

Mouse models of POD and AD feature synaptic dysfunction and loss as pathological correlates of cognitive decline [22–24]. To this end, we quantified synaptic punctae corresponding to pre-(Piccolo) and post-synaptic (Homer1, PSD95) elements in the SLM of mice 24 hours post-surgery (**Figure 4**). Surgery reduced Piccolo+ synaptic punctae by 33% (p = 0.011; **Figure 4A, C**), and Homer1+ and PSD95+ postsynaptic punctae by 10-14% (p = 0.023 and p = 0.046, respectively; **Figure 4A-B, D-E**) relative to naïve controls. URMC-099 pretreatment prevented loss of Piccolo+, PSD95+ and Homer1+ punctae comparable to control levels.

**Figure 4.**
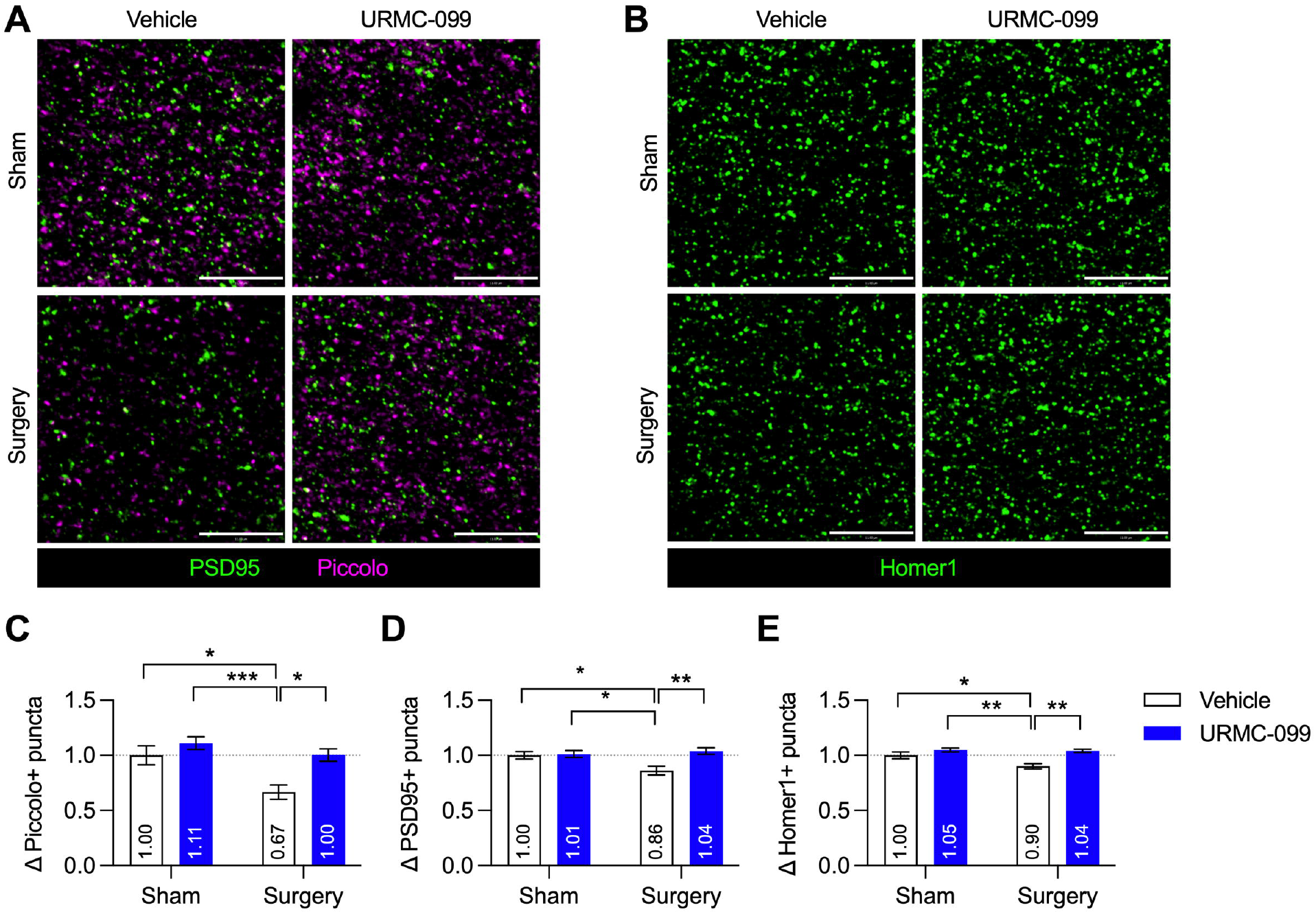
URMC-099 prevents synapse loss in the SLM of CVN-AD mice following orthopedic surgery. Six-month-old CVN-AD mice (N=6/group) received three doses i.p. of URMC-099 (10 mg/kg) prior to undergoing sham or orthopedic surgery. Brains were harvested 24 h post-surgery for IHC. (A) Representative ROIs depicting PSD95+ (postsynaptic) and Piccolo+ (presynaptic) puncta; scale bar = 11 μm. (B) Representative ROIs depicting Homer1 + (postsynaptic) puncta; scale bar = 11 μm. (C) Quantification of presynaptic Piccolo+ puncta, represented as fold change. (D) Quantification of postsynaptic PSD95+ puncta, represented as fold change. (E) Quantification of postsynaptic Homer1+ puncta, represented as fold change. Values presented as mean ± SEM (N = 6); * P < 0.05, ** P < 0.01, *** P < 0.001 for indicated comparisons; two-way ANOVA with Holm-Sidak’s multiple comparison test (C-E).

### URMC-099 prophylaxis prevents the induction of MafB in microglia following orthopedic surgery in CVN-AD mice

Lineage-specific transcription factors have recently been shown to regulate pathological microglial states under neurodegenerative and inflammatory conditions [25, 26]. In particular, the transcription factor MafB was shown recently to drive activation of spinal cord microglia in a mouse model of neuropathic pain [26]. Further, we have recently demonstrated that basal levels of MafB in microglial cells are dependent on availability of lipids in the extracellular milieu either from serum or from apoptotic debris of dying microglial cells[27]. Not only are apoptotic cells rich sources of lipids, so too are synapses [28]. Thus, we hypothesized that the neurodegenerative sequelae observed in our DSD model might also drive the upregulation of microglial MafB as a marker of microgliosis. To this end, we evaluated levels of the transcription factor MafB in SLM microglia following surgery and URMC-099 prophylaxis in CVN-AD mice. We visualized microglia using the pan-macrophage marker Iba1 and the microglia-specific marker Tmem119 [29]; only Iba1 cells co-labelled with Tmem119 were defined as microglia. Tmem119+ cells accounted for >95% of Iba1+ cells in our image set (**Supplemental Figure 1**). Orthopedic surgery increased the mean MafB IR per cell in SLM microglia (p = 0.011), an effect that was abrogated by URMC-099 prophylaxis (p = 0.046; **Figure 5A-C**). MafB induction in SLM microglia appeared homogenous in our frequency histogram analysis (**Figure 5C**). Overall, URMC-099 prevented the effect of surgery on microglial MafB induction. MafB is expressed in other CNS cell types, namely neurons [26, 30, 31]. To determine whether the induction of MafB was limited to microglia, we quantified the expression of non-microglial MafB IR (i.e., Iba1^-^ MafB objects). We observed no main effects of surgery [F(1, 20) = 0.9857; p = 0.3326] nor URMC-099 treatment [F(1, 20) = 1.266; p = 0.2738] on non-microglial MafB IR (**Supplemental Figure 2**). These results indicate the microglial MafB, but not neuronal MafB, is increased following orthopedic surgery in CVN-AD mice.

**Figure 5.**
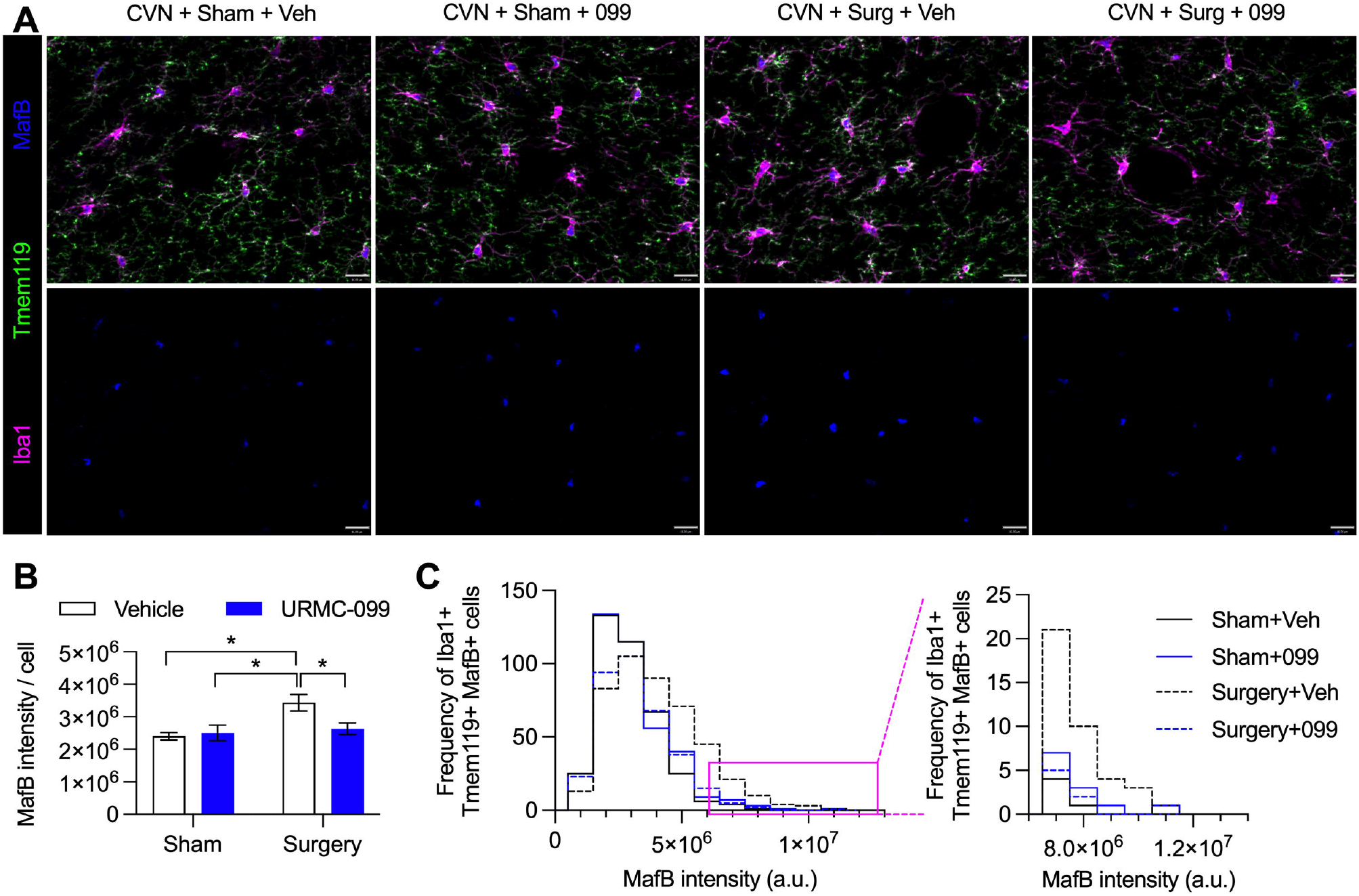
URMC-099 prevents changes in MafB microglial expression in the SLM of CVN-AD mice following orthopedic surgery. Six-month-old CVN-AD mice (N=6/group) received three doses i.p. of URMC-099 (10 mg/kg) prior to undergoing sham or orthopedic surgery. Brains were harvested 24 h post-surgery for IHC. (A) Representative images depicting the panmacrophage marker Iba1 (magenta), the microglia-specific marker Tmem119 (green), and the transcription factor MafB (blue); scale bar = 16 μm. (B) Quantification of mean MafB intensity per microglia. (C) Frequency histogram showing the number of microglia per MafB intensity bin (bin size = 1.0 × 10^6^ a. u.). (C, inset) Frequency histogram depicting cell frequencies with high MafB intensity. Values presented as mean ± SEM (N = 6); * P < 0.05 for indicated comparisons; twoway ANOVA with Holm-Sidak’s multiple comparison test (B, E).

### Orthopedic surgery and URMC-099 does not affect C1q deposition and microglial CD68/Iba1 immunoreactivity in CVN-AD mice

The opsonin C1q mediates microglia-dependent synapse loss in diverse contexts [23, 32] and is transcriptionally regulated by MafB [33]. These observations raise the possibility that microglia eliminate synapses *via* increased C1q deposition following orthopedic surgery. Although C1q expression was very robust throughout the SLM (**Figure 6A**), in agreement with previous findings by Stephan et al. [34] we observed no effects of surgery [F(1, 20) = 0.4938; p = 0.4903] nor URMC-099 treatment [F(1, 20) = 0.0280; p = 0.8689] on C1q deposition (**Figure 6B**). We next quantified the lysosomal marker CD68, whose expression is associated with activated phagocytic microglia [35] in Iba1+ cells in the SLM. However, these cells in the SLM contained high levels of CD68 regardless of condition; but neither orthopedic surgery [F(1, 20) = 0.8440; p = 0.3692] nor URMC-099 treatment [F(1, 20) = 0.8440; p = 3692] altered CD68 levels in Iba1+ cells (**Figure 6C**). We then quantified changes in microglia activation utilizing the pan-myeloid marker, Iba1. Although not well understood, during microgliosis expression of Iba1 is increased along with a thickening of processes. After surgery we found there is a positive trend of Iba1 expression along with a negative trend in surface area compared to sham (P = 0.0511 and P = 0.0921, respectively; **Figure 6C-D**). Taken together, these results demonstrate that C1q, in contrast to its roles during brain development and models of AD [23, 32], does not participate in microglial phagocytosis of synapses in this orthopedic model of DSD.

**Figure 6.**
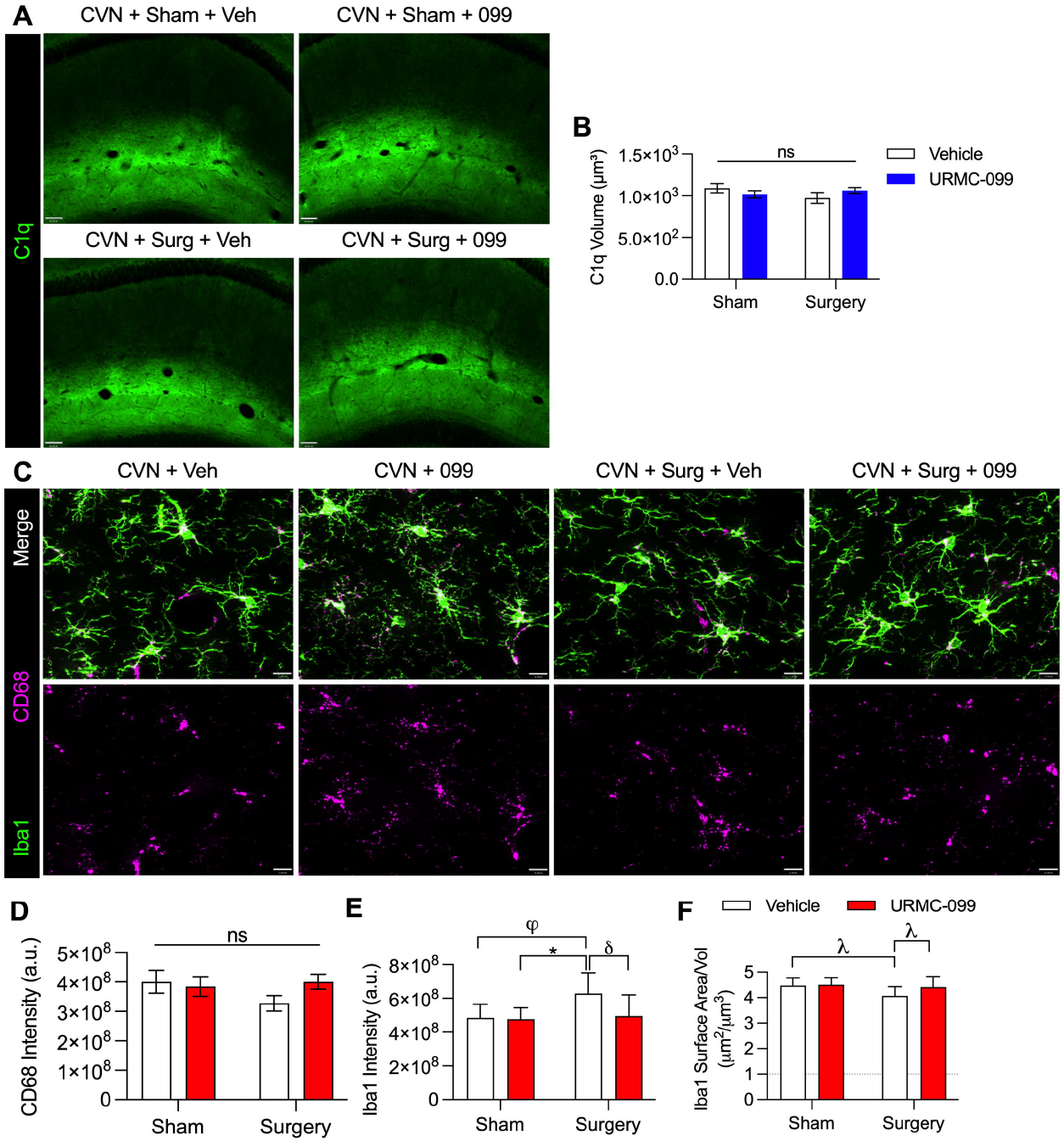
URMC-099 pretreatment does not change C1q or CD68 expression levels after surgery in the SLM of CVN-AD mice following orthopedic surgery. Six-month-old CVN-AD mice (N=6/group) received three doses i.p. of URMC-099 (10 mg/kg) prior to undergoing sham or orthopedic surgery. (A) Representative image of C1q (green) localizing in the SLM; scale bar = 60μm. (B) Quantification of C1q volume in the SLM. (C) Representative images depicting the panmacrophage marker Iba1 (green), the lysosomal marker CD68 (magenta); scale bar = 11 μm. No effects were observed on CD68 expression (D), but surgery increased Iba-1 expression and decreased surface area with a trend in URMC-099-mediated decreases (E, F). Values presented as mean ± SEM (N = 6); * p < 0.05; φ p = 0.0511; *δ* p = 0.0636; λ p = 0.0921 for indicated comparisons; two-way ANOVA with Holm-Sidak’s multiple comparison test (B-D).

Based on our results in this model of DSD, we have summarized URMC-099’s different sites of action in a visual abstract (**Figure 7**). Overall, URMC-099 pretreatment prior to surgery exerts therapeutic effects *via* inhibition of several kinase pathways in multiple vascular, immune and neuronal cell types that include preventing potential VCAM-1-mediated effects, which we hypothesize results in increased transcellular passage of inflammatory leukocytes in post-capillary venules in the SLM, deactivation of the microglial transcription factor MafB, and rescue of vulnerable pre- and postsynaptic elements in the SLM.

**Figure 7.**
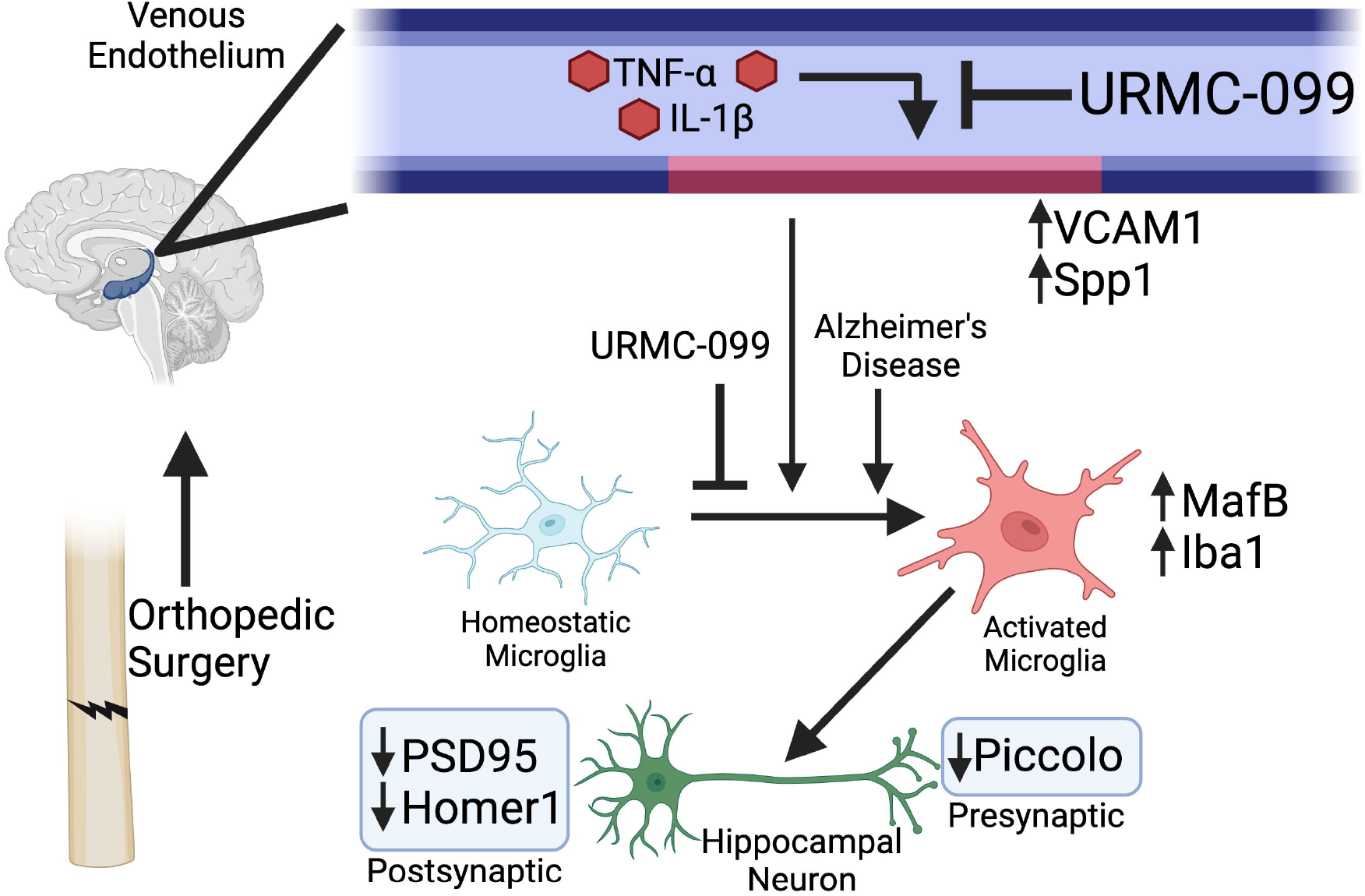
Proposed relationships between URMC-099 pretreatment and changes in endothelial, microglial and synaptic markers following surgery in six-month-old CVN-AD mice. Figure made in ©BioRender - biorender.com

## Discussion

Here, we show that URMC-099 prophylaxis is effective in the CVN-AD mouse model of DSD *prior* to the onset of age-dependent progressive vascular injury by mitigating brain endothelial cell activation (i.e., VCAM-1 and CD31 induction), vascular damage (fibrinogen leak and NET deposition), and synaptic loss. The correlation of these events with the induction of microglial MafB immunoreactivity suggests, but does not prove, a role for microglia in mediating the effects of systemic inflammation (i.e., orthopedic surgery) on synaptic integrity in these mice. Our findings are consistent with the model proposed by Yousef et al. (2019) [12] to explain the deleterious effects of aged plasma on microglia. In fact, this model posits pro-inflammatory cytokines present in the systemic milieu of aged mice (e.g., TNF-α and IL-1β) — the same ones induced systemically in our orthopedic surgery model [36, 37] — upregulate VCAM-1 on BECs, leading to sustained inflammatory activity at the BBB interface and transmission of inflammatory signals from the inflamed endothelium to microglia [12] and by extension, microglial phagocytosis of hippocampal synapses. Although it can be hypothesized that URMC-099 may exert its effects peripherally leading to cerebrovascular dysfunction, we have previously shown that URMC-099 does not affect neutrophil expansion [15]. It can still be argued that URMC-099’s therapeutic potential may act directly on modulating neutrophil activation and the release of NETs. Specifically MLK3, one of the main targets of URMC-099, is responsible for neutrophil chemotaxis [38]. Nonetheless, we contend that URMC-099 exerts at least some of its therapeutic effects by inhibiting vascular signaling through leukocyte adhesion molecules (VCAM-1 and PECAM-1/CD31) and preservation of the BBB that would otherwise initiate the inflammatory cascade that leads to microgliosis and synaptic loss.

Indeed, systemic factors induce changes in microglial reactivity by multiple mechanisms. BECs relay systemic inflammatory signals to microglia *via* soluble factors [39, 40] and direct microglial contacts with BECs [41–44]. In addition, breakdown of BEC tight junctions permits the direct entry of systemic factors (e.g., fibrinogen, systemic complement protein, etc.), many of which profoundly affect microglial function [45, 46]. Finally, peripheral leukocytes can infiltrate the CNS and release inflammatory factors to nearby cells. Indeed, CCR2+ monocytes infiltrate the hippocampal parenchyma *via* the choroid plexus following orthopedic surgery in mice [47, 48]. In the present study, the vast majority (>95%) of SLM Iba1+ cells also expressed the microglia-specific marker Tmem119 [49], indicating that they were not peripherally derived. We believe the Tmem119-negative cells were either perivascular macrophages, which lack Tmem119 expression [50], or reactive microglia that have down-regulated their expression of this marker [51]. In addition, we did not observe any Ly6G+ neutrophils in the brain parenchyma. However, the selective increase of VCAM-1 on post-capillary venules, through which leukocytes preferentially transmigrate [52], supports the possibility of increased tethering of leukocytes to the endothelium and infiltration in the SLM following orthopedic surgery, although such infiltration may be limited to the perivascular space rather than the CNS parenchyma.

In our model, orthopedic surgery induced upregulation of VCAM-1 in SLM postcapillary venules (**Figure 2–3**), damage to the BBB **(Figure 3)**, loss of synaptic elements (**Figure 4**), and increased MafB immunoreactivity in SLM microglia (**Figure 5**). Substrates that may drive induction of MafB following surgery are not well characterized. Both oxidative stress [53] and engulfment of apoptotic cells [33] have been shown to induce MafB expression in macrophages. Because efferocytosis is dependent on LXR [54], it is conceivable that orthopedic surgery, when combined with pre-existing neurodegeneration (i.e., AD pathology), causes microglial engulfment of focal apoptotic synapses (i.e., synaptosis) and other cellular debris [55], leading to MafB induction. A recent report showing microglial induction of MafB following peripheral nerve injury supports this hypothesis [26].

Microglia pathologically interact with synaptic elements in mouse models of neurodegeneration and neuroinflammation [23, 42, 56–58]. Because microglia mediate synapse loss in mouse models of AD *via* complement-dependent pruning processes [23], and tibial fracture/fixation upregulated MafB, which in turn transcriptionally regulates C1q [33], we investigated whether synapse loss in our DSD model might also depend on C1q. Here we did not observe changes in C1q deposition or microglial CD68 immunoreactivity following orthopedic surgery in CVN-AD mice (**Figure 6**). It is worth noting, however, that C1q is deposited at high levels in the SLM of wild-type [34] and CVN-AD mice (this study). Moreover, we observed high levels of CD68 selectively in SLM microglia compared to surrounding regions across all our conditions. Thus, these two factors may predispose the SLM of CVN-AD mice to synaptic loss following systemic and/or CNS inflammation. We cannot discount the possibility that synaptic loss may be due to fibrinogen leakage. In the context of AD, fibrinogen has been shown to bind to CD11b causing synaptic pruning independent of amyloid plaques [46]. Finally, we cannot completely exclude the possibility that our synaptic data reflects changes in the expression of genes encoding Piccolo, PSD95, and Homer1, as has been observed in AD patients [59], or the clustering of these proteins in synaptic structures (that would influence their ability to form punctae). Even so, changes in synaptic gene expression correlate with cognitive impairment in AD patients [60, 61], and microglial depletion ameliorates pathological changes in synaptic gene expression in mouse AD models [62]. Taken together, we believe the amelioration of synaptic loss we observed in our model of DSD is likely microglial-dependent and these are tractable targets for URMC-099 prophylaxis in protecting the neurovascular unit from postoperative complications including delirium and dementia. Overall, using our orthopedic model of tibial fracture/fixation in six-month-old CVN mice, we have demonstrated that damage to the BBB resulting in endothelial activation with increased expression of VCAM-1 in post-capillary venules of hippocampal SLM, associated microgliosis and damage to synaptic architecture, can be prevented with prophylactic administration of the MLK3/LRRK2 broad-spectrum inhibitor, URMC-099. Based on these findings, we conclude that this therapeutic approach may represent an intervention to significantly decrease deleterious effects of DSD in the elderly population at risk for morbidity and mortality after surgery.

## Declarations

All manuscripts must contain the following sections under the heading ‘Declarations’:

### • Ethics approval

The procedures followed to provide tissues from APPSwDI/mNos2^-/-^ (CVN-AD) mice were performed in strict compliance with animal protocols approved by the Institutional Animal Care and Use Committees (IACUC) of Duke University (A249-17-11).

### • Consent for publication

All authors have reviewed the data and its interpretation and consent to the submission of this publication.

### • Availability of data and material

All data generated or analyzed during this study are included in this published article and its supplementary information files.

### • Authors’ contributions

PMR performed the experiments, analyzed the data and wrote and edited the manuscript. HL performed the experiments, analyzed the data, and wrote and edited the manuscript. RV performed experiments on CVN-AD mice. NT and HAG. supervised PMR and HL and contributed to the interpretation of the data.

## Acknowledgements

We thank Angela Stout in the Gelbard lab for her technical expertise.

## • Authors’ information (optional)

See front page

---

**Supplement Figure 1**. Iba1+ cells in the SLM are predominantly microglia. Six-month-old CVN mice (N=6/group) received three doses i.p. of URMC-099 (10 mg/kg) prior to undergoing sham or orthopedic surgery. Brains were harvested 24 h post-surgery for IHC. (A) Quantification of Tmem119hi and Tmem119lo Iba1+ cells in the SLM for surgical and URMC-099 treatment groups. (B) Quantification of mean Tmem119hi and Tmem119lo cell numbers per 40X field of view. Values presented as mean ± SEM (n = 6); two-way ANOVA with Holm-Sidak’s multiple comparison test (B).

**Supplement Figure 2**. Orthopedic surgery and URMC-099 treatment had no effect on non-microglial MafB immunoreactivity in the SLM. Six-month-old CVN-AD mice (N=6/group) received three doses i.p. of URMC-099 (10 mg/kg) prior to undergoing sham or orthopedic surgery. Brains were harvested 24 h post-surgery for IHC. Representative image of MafB (magenta) co-labeling oligodendrocyte-specific marker (CNP1, green), neuronal-specific marker (NeuN, green), and astrocyte-specific marker (GFAP, green) (A, B, and C respectively); scale bar = 16 μm (A, B) 11 μm (C). (D) Frequency histogram showing the number of microglia per MafB intensity bin (bin size = 1.0 × 106 a. u.). (D) Quantification of mean MafB intensity normalized to the number of Iba1-MafB+ nuclei. Values presented as mean ± SEM (n = 6); two-way ANOVA with Holm-Sidak’s multiple comparison test (D).

