## Supplementary figures for "URMC-099 Prophylaxis Prevents Hippocampal Vascular Vulnerability and Synaptic Damage in an Orthopedic Model of Delirium Superimposed on Dementia"

**Figure 1. URMC-099 inhibits BEC activation *in vitro***. Serum-deprived Bend.3 cells were pre-treated with URMC-099 (099) or DMSO for 30 min, after which cells were stimulated with IL-1β (10 ng/mL). (A) Representative images of VCAM-1 and PECAM-1 staining in bend.3 cells; scale bar = 15 μm. (B) Quantification of PECAM-1 IR, represented as fold change. (C) Quantification of VCAM-1 IR, represented as fold change. (D) qPCR analysis of bend.3 transcriptional response to IL-1B and URMC-099 treatment. Values presented as mean ± SEM (N = 4 cell culture wells). ** P < 0.01, *** P < 0.001, **** P < 0.0001 for indicated comparisons; one-way ANOVA with Dunnett’s multiple comparison test (B-D).

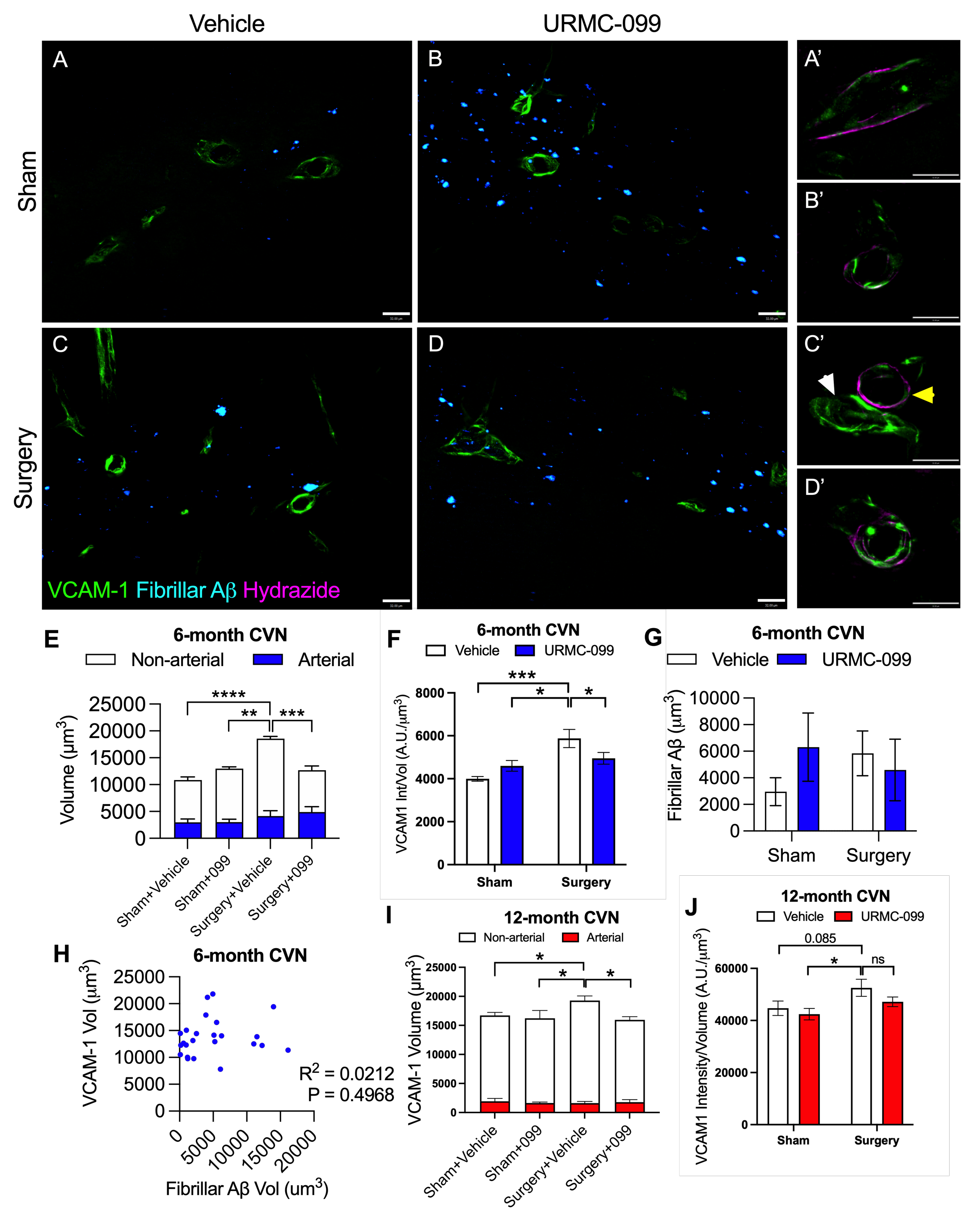

**Figure 2. URMC-099 prophylaxis prevents the induction of VCAM-1 in the SLM of 6-month-old CVN-AD mice following surgery.** Six and twelve-month-old CVN-AD mice (N=6/group) received three doses i.p. of URMC-099 (10 mg/kg) prior to undergoing sham or orthopedic surgery. Brains were harvested 24 h post-surgery for IHC. (A-D) Representative images depicting VCAM-1 (green) and fibrillar AB (cyan). (A’-D’) Representative images depicting arterial VCAM-1 vessels using the artery-specific dye Alexa 633-hydrazide (magenta); images correspond to experimental conditions represented by A-D; scale bar = 32μm. (E) Quantification of total VCAM-1 volume (full bars), venous VCAM- 1 volume (white bars), and arterial VCAM-1 volume (blue bars) at 6-months age; statistical comparisons are shown for total VCAM-1. (F) Quantification of VCAM-1 intensity normalized to volume at 6-months of age. (G) Quantification of fibrillar AB. (H) Correlation between total VCAM-1 volume and fibrillar AB volume at 6-months of age. (I) Quantification of total VCAM-1 volume (full bars), venous VCAM- 1 volume (white bars), and arterial VCAM-1 volume (blue bars) at 12-months age; statistical comparisons are shown for total VCAM-1. (J) Quantification of VCAM-1 intensity normalized to volume at 12-months of age. Values presented as mean ± SEM (N = 6); * P<0.05, ** P< 0.01, *** P < 0.001, **** P < 0.0001 for indicated comparisons; two-way ANOVA with Holm-Sidak’s multiple comparison test (E, F) or Pearson’s coefficient (D)

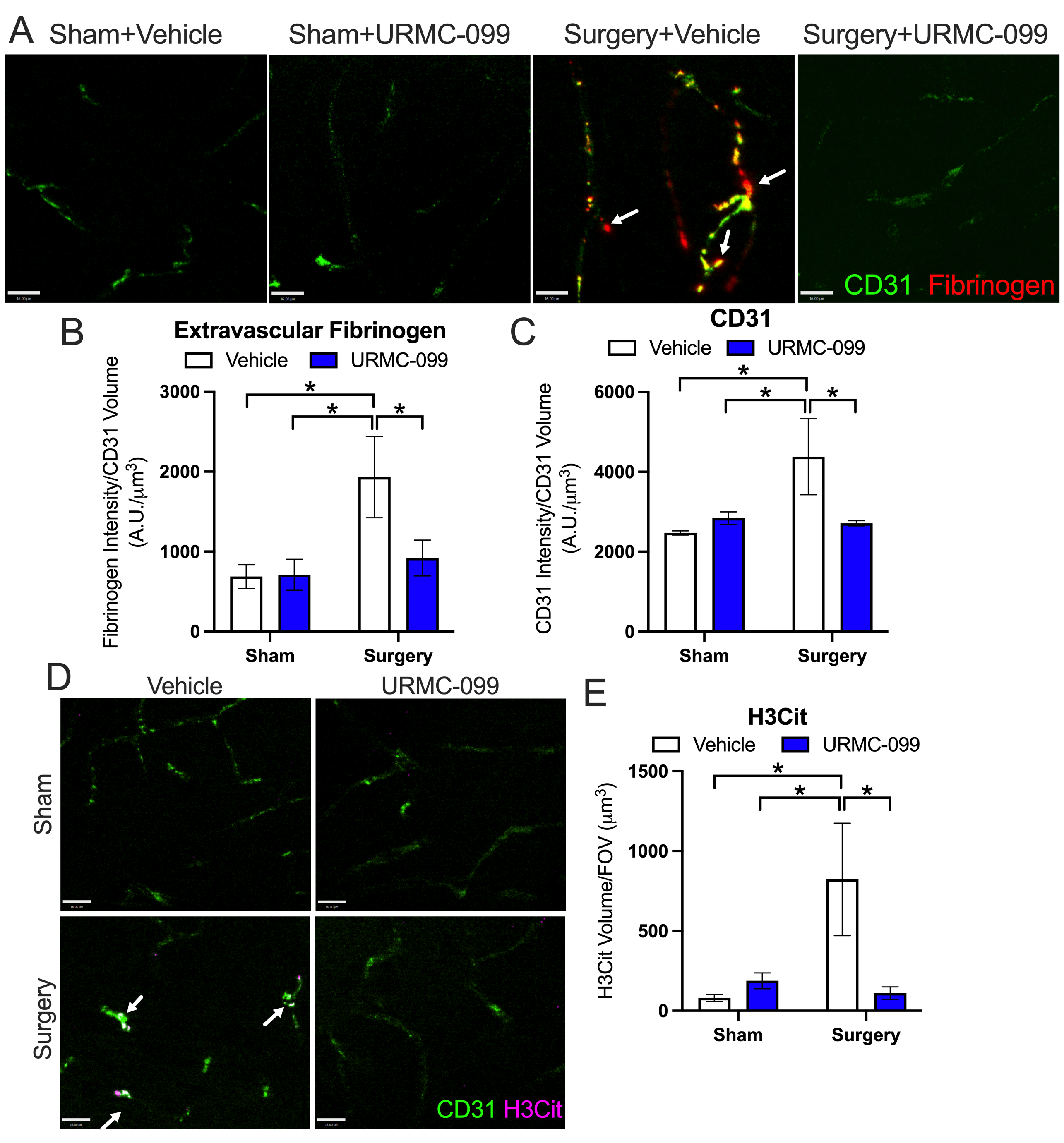

**Figure 3. URMC-099 prevents vascular damage in the SLM of 6-month CVN-AD mice following orthopedic surgery**. Six-month-old CVN-AD mice (N=6/group) received three doses i.p. of URMC-099 (10 mg/kg) prior to undergoing sham or orthopedic surgery. Brains were harvested 24 h post-surgery for IHC. (A) Representative images depicting CD31/Pecam1 (green) and fibrinogen (red); scale bar = 16 μm. (B) Quantification of extravascular fibrinogen. (C) Quantification of CD31/Pecam1 intensity normalized to CD31/Pecam1 volume. (D) Representative images depicting CD31/Pecam1 (green) and NET marker, H3Cit (magenta); scale bar = 16 μm. (E) Quantification of neutrophil NET marker, H3Cit. Values presented as mean ± SEM (N = 6); * P < 0.05 for indicated comparisons; two-way ANOVA with Holm-Sidak’s multiple comparison test (C-E).

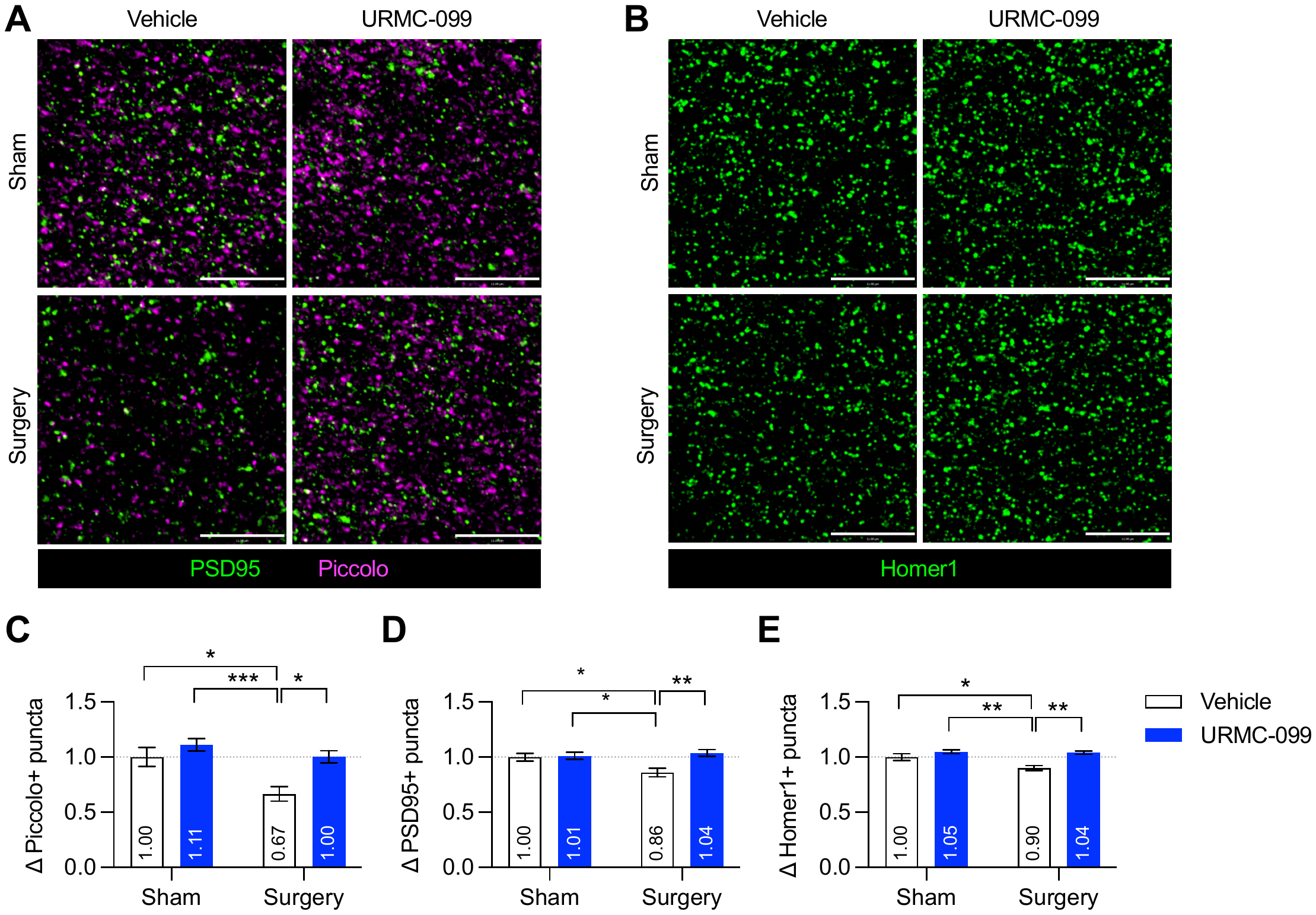

**Figure 4. URMC-099 prevents synapse loss in the SLM of CVN-AD mice following orthopedic surgery**. Six-month-old CVN-AD mice (N=6/group) received three doses i.p. of URMC-099 (10 mg/kg) prior to undergoing sham or orthopedic surgery. Brains were harvested 24 h post-surgery for IHC. (A) Representative ROIs depicting PSD95+ (postsynaptic) and Piccolo+ (presynaptic) puncta; scale bar = 11 μm. (B) Representative ROIs depicting Homer1+ (postsynaptic) puncta; scale bar = 11 μm. (C) Quantification of presynaptic Piccolo+ puncta, represented as fold change. (D) Quantification of postsynaptic PSD95+ puncta, represented as fold change. (E) Quantification of postsynaptic Homer1+ puncta, represented as fold change. Values presented as mean ± SEM (N = 6); * P < 0.05, ** P < 0.01, *** P < 0.001 for indicated comparisons; two-way ANOVA with Holm-Sidak’s multiple comparison test (C-E).

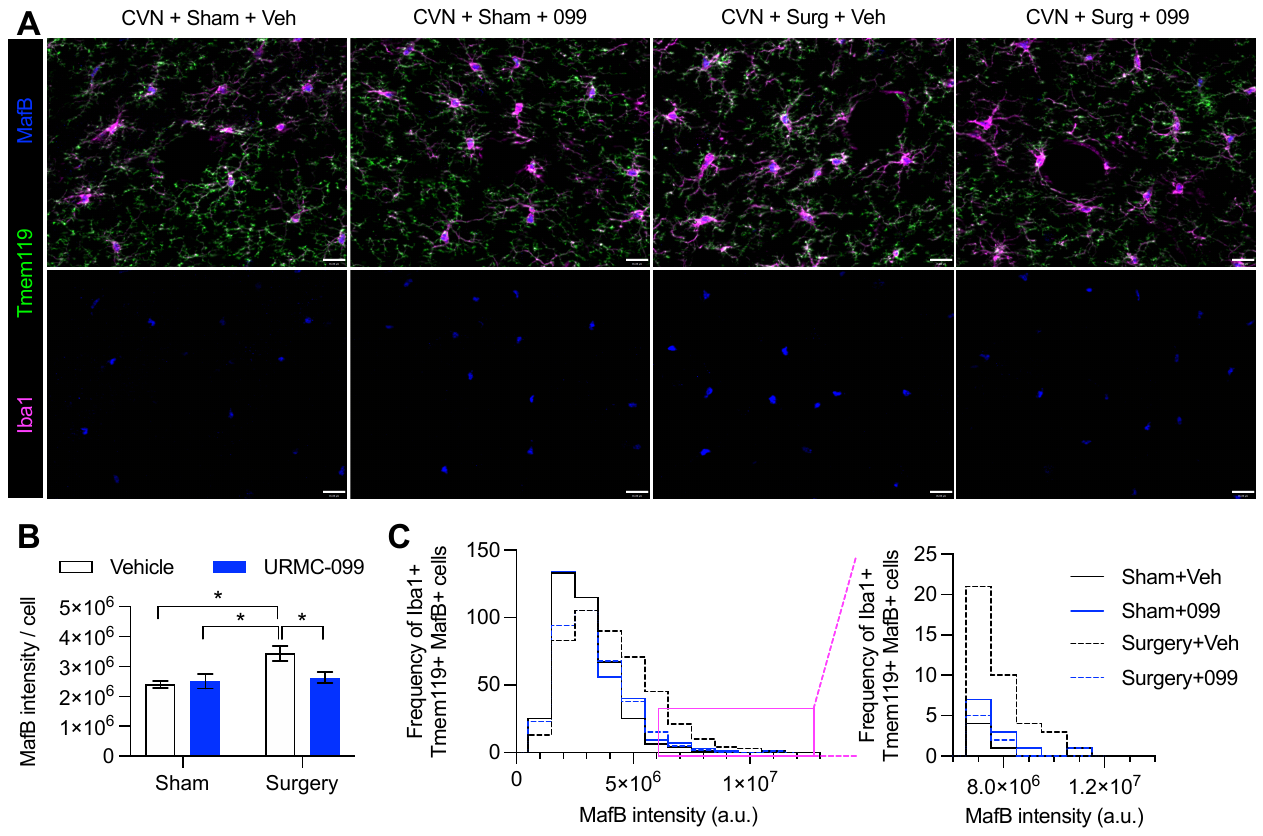

**Figure 5. URMC-099 prevents changes in MafB microglial expression in the SLM of CVN-AD mice following orthopedic surgery.** Six-month-old CVN-AD mice (N=6/group) received three doses i.p. of URMC-099 (10 mg/kg) prior to undergoing sham or orthopedic surgery. Brains were harvested 24 h post-surgery for IHC. (A) Representative images depicting the pan-macrophage marker Iba1 (magenta), the microglia-specific marker Tmem119 (green), and the transcription factor MafB (blue); scale bar = 16 μm. (B) Quantification of mean MafB intensity per microglia. (C) Frequency histogram showing the number of microglia per MafB intensity bin (bin size = 1.0 ⨉ 10^6^ a. u.). (C, inset) Frequency histogram depicting cell frequencies with high MafB intensity. Values presented as mean ± SEM (N = 6); * P < 0.05 for indicated comparisons; two-way ANOVA with Holm-Sidak’s multiple comparison test (B, E).

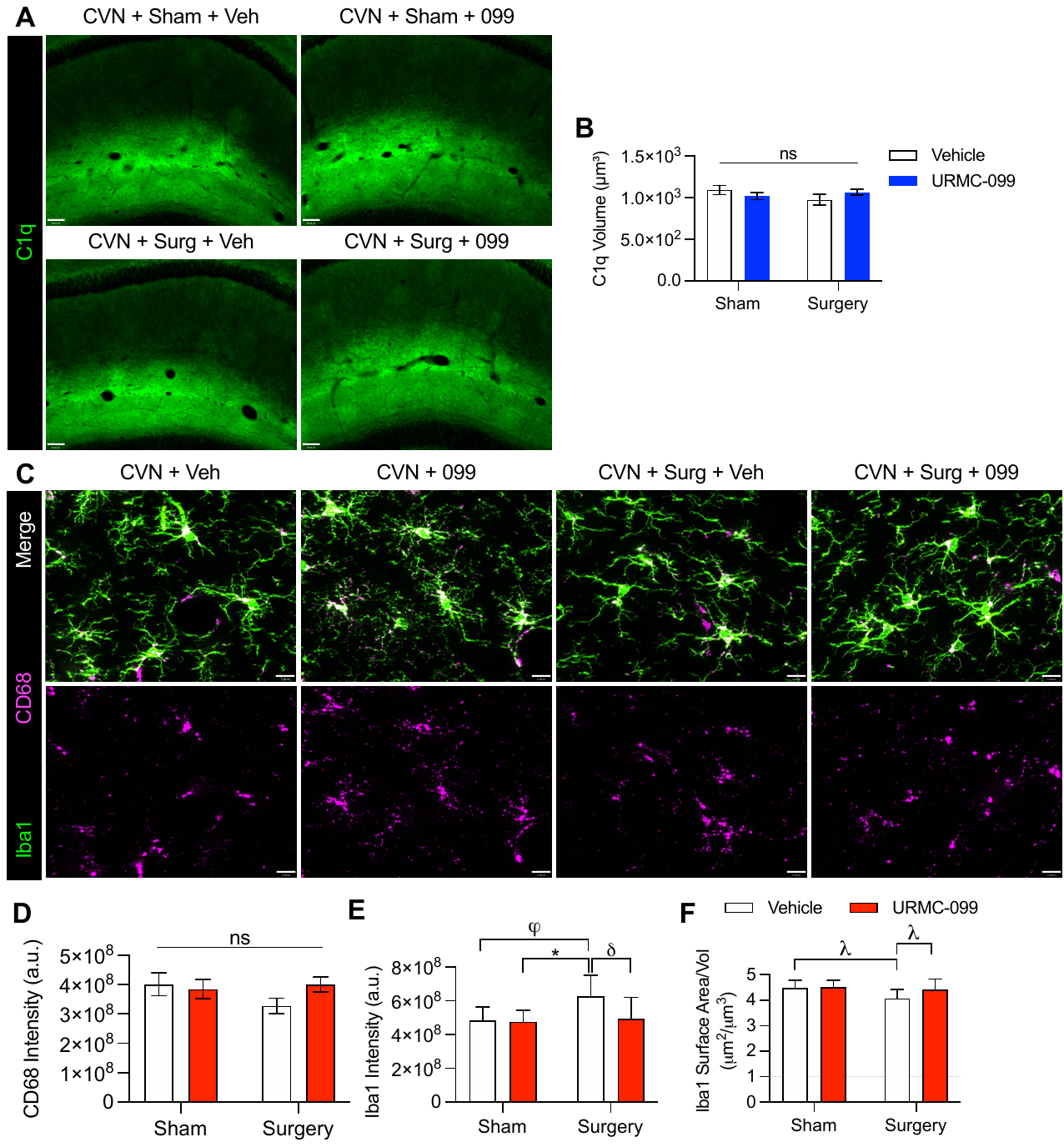

**Figure 6. URMC-099 pretreatment does not change C1q or CD68 expression levels after surgery in the SLM of CVN-AD mice following orthopedic surgery.** Six-month-old CVN-AD mice (N=6/group) received three doses i.p. of URMC-099 (10 mg/kg) prior to undergoing sham or orthopedic surgery. (A) Representative image of C1q (green) localizing in the SLM; scale bar = 60μm. (B) Quantification of C1q volume in the SLM. (C) Representative images depicting the pan-macrophage marker Iba1 (green), the lysosomal marker CD68 (magenta); scale bar = 11 μm. No effects were observed on CD68 expression (D), but surgery increased Iba-1 expression and decreased surface area with a trend in URMC-099-mediated decreases (E, F). Values presented as mean ± SEM (N = 6); * p < 0.05; φ p = 0.0511; 𝛿 p = 0.0636; λ p = 0.0921 for indicated comparisons; two-way ANOVA with Holm-Sidak’s multiple comparison test (B-D).

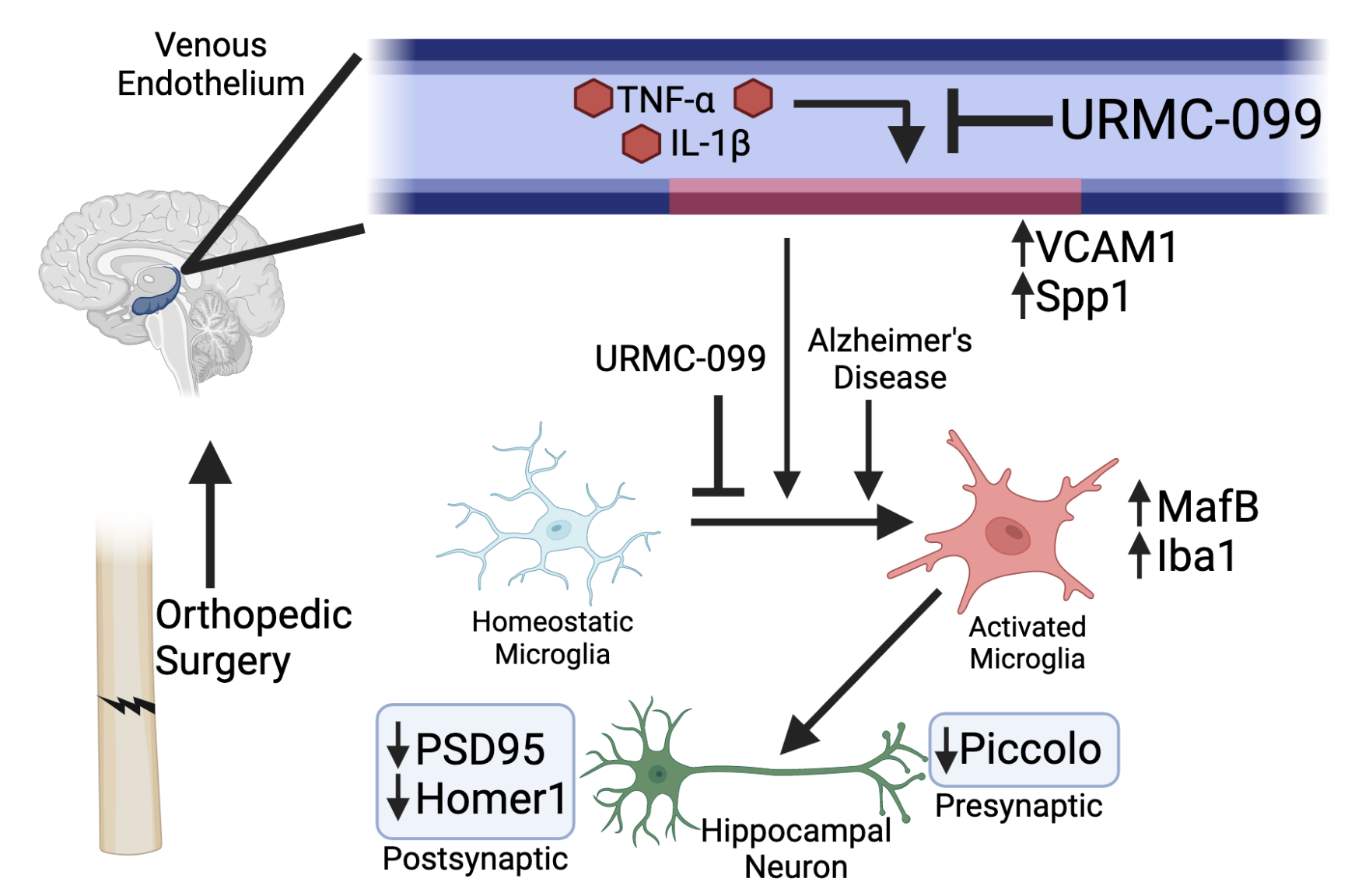

**Figure 7.** Proposed relationships between URMC-099 pretreatment and changes in endothelial, microglial and synaptic markers following surgery in six-month-old CVN-AD mice.

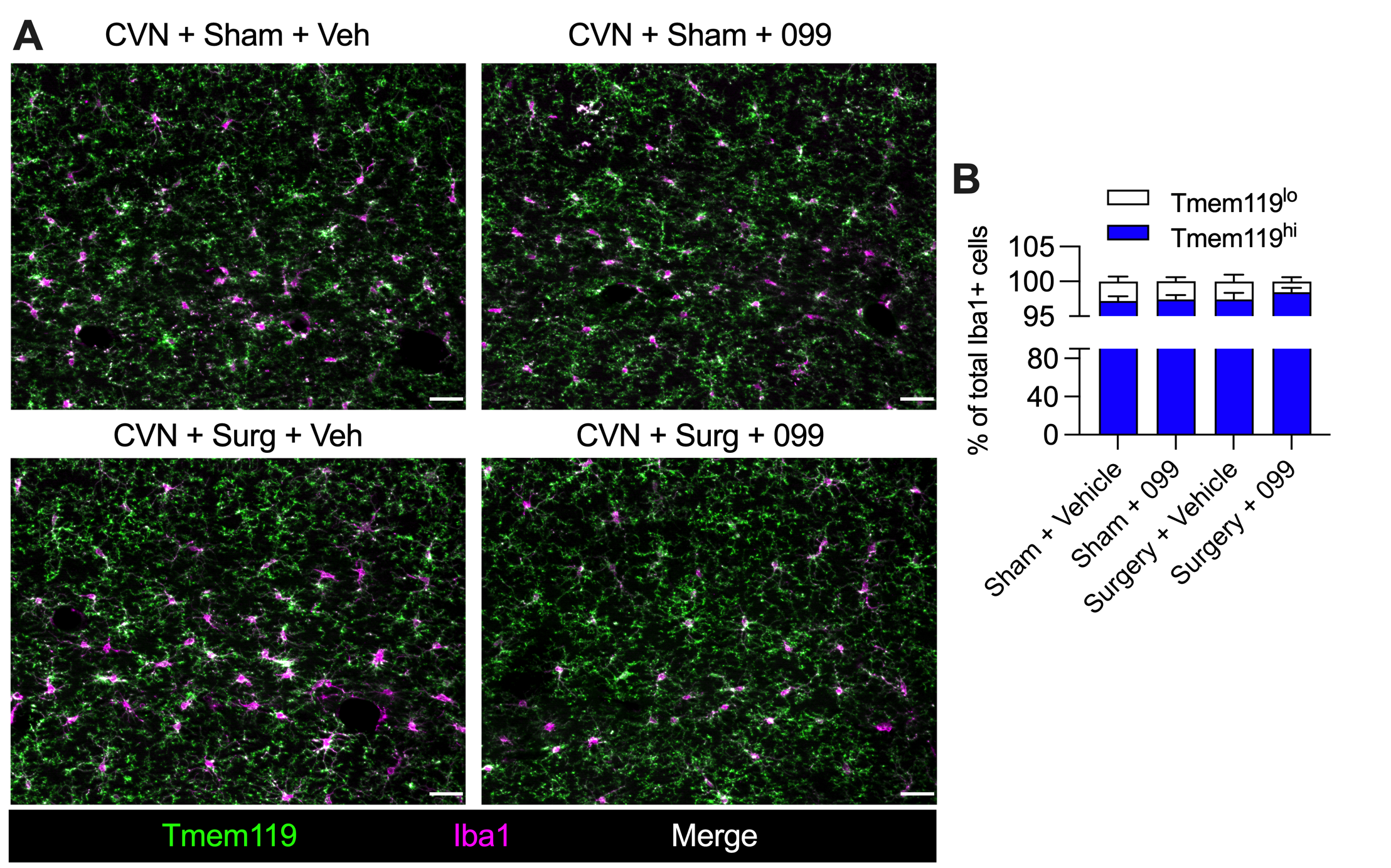

**Supplement Figure 1.** Iba1+ cells in the SLM are predominantly microglia. Six- month-old CVN mice (N=6/group) received three doses i.p. of URMC-099 (10 mg/kg) prior to undergoing sham or orthopedic surgery. Brains were harvested 24 h post-surgery for IHC. (A) Quantification of Tmem119^hi^ and Tmem119^lo^ Iba1+ cells in the SLM for surgical and URMC-099 treatment groups. (B) Quantification of mean Tmem119^hi^ and Tmem119^lo^ cell numbers per 40X field of view. Values presented as mean ± SEM (n = 6); two-way ANOVA with Holm-Sidak’s multiple comparison test (B).

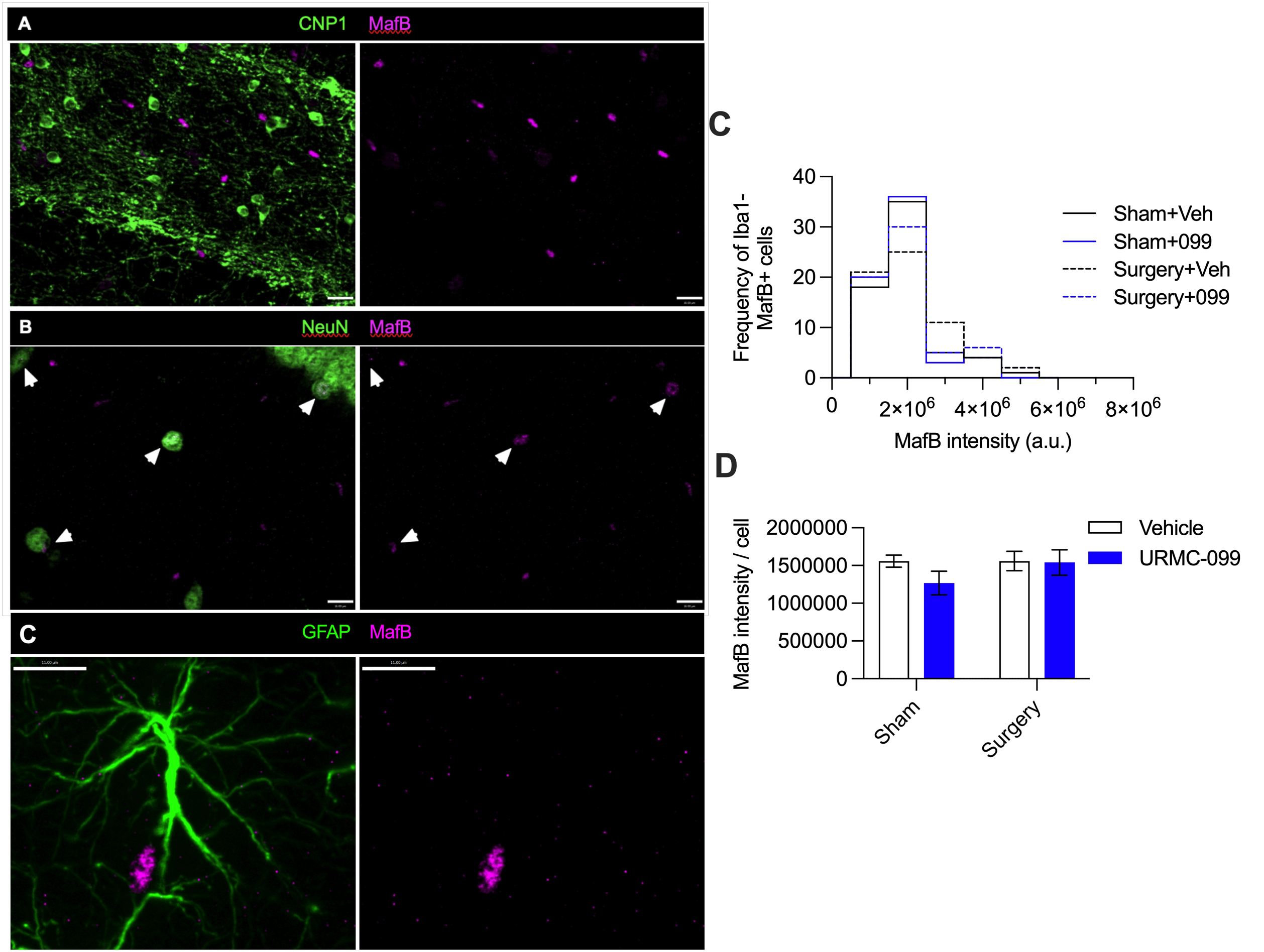

**Supplement Figure 2.** Orthopedic surgery and URMC-099 treatment had no effect on non-microglial MafB immunoreactivity in the SLM. Six-month-old CVN-AD mice (N=6/group) received three doses i.p. of URMC-099 (10 mg/kg) prior to undergoing sham or orthopedic surgery. Brains were harvested 24 h post-surgery for IHC. Representative image of MafB (magenta) co-labeling oligodendrocyte-specific marker (CNP1, green), neuronal-specific marker (NeuN, green), and astrocyte-specific marker (GFAP, green) (A, B, and C respectively); scale bar = 16 μm (A, B) 11 μm (C). (D) Frequency histogram showing the number of microglia per MafB intensity bin (bin size = 1.0 ⨉ 10^6^ a. u.). (D) Quantification of mean MafB intensity normalized to the number of Iba1^-^ MafB^+^ nuclei. Values presented as mean ± SEM (n = 6); two-way ANOVA with Holm-Sidak’s multiple comparison test (D).
